# Whole genome assembly and annotation of the Bumblebee Wax Moth, *Aphomia sociella*

**DOI:** 10.1101/2025.06.18.660359

**Authors:** Ronja Marlonsdotter Sandholm, Gustav Vaaje-Kolstad, Sabina Leanti La Rosa

**Affiliations:** Faculty of Chemistry, Biotechnology and Food Science, NMBU - Norwegian University of Life Sciences, Ås, Norway

## Abstract

The bumble bee wax moth, *Apshomia sociella*, is an important lepidopteran pest impacting bee colonies essential for pollination and apiculture. Despite its relevance, sequence efforts aimed at understanding the genetic makeup of this species have not yet been undertaken. In this work, we successfully achieved a high-quality de novo genome assembly of *A. sociella* and comprehensive gene annotations generated from long-read DNA and RNA sequencing with Oxford Nanopore technology. The haploid assembly includes 347 contigs, with an N50 of 4.96 Mb, and contains 27 242 protein-coding genes. Benchmarking Universal Single Copy Orthologs (BUSCO) analyses indicates that the assembly has a high level of completeness (98.3%) and low level of fragmentation (0.6%) and duplication (3.9%). Phylogenomic analyses with other members of the Lepidoptera order placed *A. sociella* in the same clade as *Corcyra cephalonica* and indicates close evolutionary relationships with the other two species in the subfamily *Galleriinae*, namely *Achroia grisella* and *Galleria mellonella*. This new high-quality genome assembly represents a valuable resource for investigating the genomic basis of ecological specialization of this species and offers critical support for research aimed at developing sustainable and effective pest management strategies.

## Introduction

In recent years, sequencing the genomes of all living species has emerged as a cornerstone goal in biodiversity science. Initiatives like the Earth BioGenome Project aim to generate high-quality reference genomes for all eukaryotic life on Earth, creating a comprehensive molecular archive of Earth’s biological diversity (Lewin et al., 2022). These reference genomes serve as foundational resources that empower a wide range of scientific and practical applications, including studies of evolution and gene function, conservation genetics and species breeding programs. Moreover, for species that pose serious economic threats to agriculture and farming, elucidating the underlying genetic determinants can facilitate the development of novel, targeted control strategies.

The order Lepidoptera consists of butterflies and moths, with around 160,000 described species (van Nieukerken et al., 2011). This makes the order one of the largest in the animal kingdom, second only to Coleoptera (also known as beetles). Lepidoptera includes the family *Pyralidae*, of which the wax moths belong. To date, 28 out of 6440 species in this family have publicly available genomes (Khramov, 2025; Schoch et al., 2020). This family of moths is a diverse group, with most members being inconspicuous. However, many members are economically important pests, infesting key agricultural products, such as dried fruits and vegetables, grains and seeds. Within this family, the subfamily *Galleriinae* includes notable species such as the bumblebee wax moth *Aphomia sociella, the* greater wax moth *Galleria mellonella* and the lesser wax moth *Achroia grisella*. While wax moth larvae are pests of beehives, *A. sociella* primarily attacks bumblebee nests but can also invade weakened commercial honeybee colonies (Krams et al., 2025). Females deposit their eggs within the nests, where the larvae feed on wax comb and honey, causing severe damage that can lead to colony collapse. Given that bumblebees play a crucial commercial role in agriculture by providing essential pollination services for many crops across temperate regions of the Northern Hemisphere, moth infestations pose significant environmental threats and economic challenges that profoundly impact beekeeping sustainability.

In addition to their role as pests, several moths, including *A. sociella*, have been reported to consume plastics. In recent years, *G. mellonella* and *A. grisella* have gained particular interest for its “plastivore” lifestyle and has become increasingly studied as a potential source of plastic degrading enzymes, allegedly originating both from the insect itself and from its endogenous gut microbiome (Ren et al., 2019; Zhang et al., 2020). Despite interest in the plastic degrading abilities of insect larvae, the evidence remains controversial, as studies have reported inconsistent results regarding plastic oxidation or bioassimilation (Réjasse et al., 2022; Stepnov et al., 2024).

Control strategies for managing moth populations have included the use of pheromone-baited traps and bioinsecticides composed of lethal viral or bacterial pathogens targeting the larval stages (Niu et al., 2024; Zhang et al., 2021). Pheromones are release by both males and females to attract each other during mating (Kindl et al., 2012; Kindl et al., 2011). Targeting pheromones offers an effective approach for selective pest control is due to their species specificity, high potency in very small amounts, and low toxicity to non-target animals (Witzgall et al., 2010). While there is a limited understanding of the identity, genetic basis, and regulation of pheromone production in *A. sociella, s*uch knowledge could be applied to enhance the efficacy of pheromone-baited traps and bioinsecticides to control the bumblebee wax moth populations.

Here, capitalizing on advancements in long-read sequencing technology, we present a high-quality genome assembly of *A. sociella*. We obtained gene annotations through short- and long-read RNA sequencing data and demonstrated the utility of our new genome assembly through comparative genomics. We envisage that these genome resources will facilitate functional studies into the moth, giving a deeper understanding of basic biological mechanisms, such as life cycle and survival, pheromone signaling and immunity, that may be of value for developing pest control measures.

## Methods

### Tissue collection

*A. sociella* larvae were collected from a private breeder of tree bumblebees (*Bombus hypnorum*) in Enger, Innlandet county, Norway. The collection occurred on the 16^th^ of September 2022. The larvae were fed remnants of the infested bumblebee nest until they arrived in the laboratory. Note that the sex of the specimen could not be determined at the larval stage. Larvae were rinsed in sterile MilliQ water and either stored at -20 °C before DNA extraction or preserved in RNA*later*™ Stabilization solution (Invitrogen, Cat. no.: AM7020) at -80 °C before RNA extraction.

### DNA extraction and whole genome sequencing

High molecular-weight (HMW) DNA from an individual larva was extracted using a previously established protocol, with minor modifications (Mugford et al., 2020). The frozen larva was briefly minced on ice before homogenization in 750 µL cold phosphate-buffered saline (PBS, Sigma-Aldrich, Cat. No.: P4417) using a TissueRuptor II (QIAGEN, ID: 9002755), with a disposable probe (QIAGEN, ID: 990890) at 35 000 rpm for 10 seconds. The homogenized tissue was transferred to a 2 mL Eppendorf tube and centrifuged at 3 000 x g for 5 minutes at 4 °C. The resulting pellet was washed with 1 mL of cold PBS. The cetrimonium bromide:chloroform method was used to obtain HMW DNA, followed by DNA precipitation using isopropanol. The DNA was quantified using the Qubit Broad Range DNA assay (Invitrogen), according to the manufacturer’s protocol, on a Qubit 1.0 fluorometer. DNA quality was assessed using a Nanodrop One UV-Vis spectrophotometer (Thermo Scientific, Wilmington, USA). The sequencing library was prepared using the Ligation Sequencing kit SKQ-LSK110 (Oxford Nanopore Technologies) following the manufacturer’s instructions. The library was loaded for sequencing on a single R9.4.1 flow cell on a PromethION platform (Oxford Nanopore Technologies) at the Norwegian University of Life Sciences. The system was operated with the MinKNOW v22.10.5 software.

### RNA extraction and sequencing

A whole single larva of *A. sociella* was used for RNA extraction. RNA was obtained by homogenizing the tissue in 750 µL PBS with a TissueRuptor II (QIAGEN, ID: 9002755) with a disposable probe (QIAGEN, ID: 990890) at 35 000 rpm for 1 minute, followed by extraction with the RNeasy Mini Kit (QIAGEN, ID: 74104) according to the manufacturer’s instructions. RNA concentration and purity were determined using a Qubit 3.0 fluorometer and a Nanodrop One instrument (Thermo Scientific, Wilmington, USA). RNA integrity was verified using an Agilent 4200 Tape Station System (Agilent Technologies, Santa Clara, CA, USA). PolyA-enriched RNA was sequenced on the Illumina NovaSeq 6000 platform, utilizing a pair-end 300 bp sequencing strategy. Per the manufacturer’s protocol, Novogene (Beijing, China) carried out library preparations using a TruSeq Stranded mRNA kit (Illumina, San Diego, CA, USA). A long-read sequencing library of the extracted RNA was prepared using the PCR-cDNA Barcoding kit SQK-PCB111.24 (Oxford Nanopore Technologies) and sequenced on a R10.4 flow cell on a PromethION platform (Oxford Nanopore Technologies) at the Norwegian University of Life Sciences. The system was operated with the MinKNOW v23.07.12 and basecalling was performed using GPU-enabled Guppy v7.1.4 using the high-accuracy model.

### Genome assembly

Basecalling on the ONT raw reads was performed using GPU-enabled Guppy v6.3.8 with the high-accuracy model (Oxford Nanopore Technologies, 2023). Reads were filtered by quality using fastp v0.23.4 with the `--length_required 50, --qualified_quality_phred 7`and `--trim_front1 400`parameters (Chen et al., 2018). Four genome assemblies were generated using Canu v2.2 using the `-nanopore-raw`parameter (Koren et al., 2017), where one was obtained using standard parameters and three by adjusting the error rate for each assembly (`correctedErrorRate=0.01`, `correctedErrorRate=0.02,`and `correctedErrorRate=0.03`). Error correction was performed using Inspector v1.0.2 (Chen et al., 2021). Quality assessment of the assemblies was conducted using Merqury v1.3 (Rhie et al., 2020) and BUSCO v5.8.3 with the `-m geno`and `-l lepidoptera_odb12`parameters (Manni et al., 2021) . The Canu assembly with an adjusted error rate of 2% was chosen for downstream processing. Purge Haplotigs v1.1.2 (Roach et al., 2018) was used to generate a primary assembly, identify haplotigs and artefactual contigs, and eventually obtain a deduplicated, non-redundant genome assembly for the insect, with the parameters for purge_haplotigs cov being `-l 5`, `-m 100`and `-h 200`. The genome assembly was also screened for contaminations using FCS-GX v0.5.0 (Astashyn et al., 2023), and assessed using assembly_stats v0.1.4 (Trizna, 2020), both with standard parameters.

### Genome annotation

Before annotation, repeat elements of the genomes were identified and masked using RepeatModeler v4.0 (Smit & Hubley, 2008-2015) with `-database`function followed by RepeatMasker v1.0 (Smit et al., 2013-2015) using default setting for de novo prediction. Protein-coding genes were predicted using transcriptomics-based methods with short and long RNA reads. Short-read RNA sequences were filtered using FastQC v0.11.9 (Andrews, 2010) with standard parameters. A *de novo* hybrid transcriptome assembly was generated using rnaSPAdes v3.15.5 (Bushmanova et al., 2019), and the software was run with standard parameters using the short- and long-read RNA-seq data generated in this study. Funannotate v1.8.15 (Palmer, 2016) was first trained using the short-read RNA sequences and the hybrid transcriptome assembled reads from rnaSPAdes, followed by gene prediction and refining using standard settings. Predicted genes were functionally annotated with Funannotate using the Pfam (Mistry et al., 2020), InterPro (Jones et al., 2014), eggNOG (Cantalapiedra et al., 2021), UniProt (Consortium, 2022), MEROPS (Rawlings et al., 2017), CAZyme (Drula et al., 2021) and GO ontology (Ashburner et al., 2000; Consortium et al., 2023) databases. The parameters `--busco_db lepidoptera_odb10`and `--signalp`were added to include BUSCO models and secreted proteins with SignalP (Nielsen et al., 2019). KEGG Orthologs (KOs) were assigned to the annotated genes using KofamKOALA (Aramaki et al., 2020), and pathways were identified using KEGG Mapper Reconstruct (Kanehisa & Sato, 2020).

### Phylogenomic analysis

To investigate the phylogenomic relationship between *A. sociella* (this study) and other members of the Lepidoptera order, publicly available genomes were obtained and BUSCO orthologous genes (lepidoptera_odb12) aligned using OrthoFinder v3.0.1b1 (Emms & Kelly, 2019). The final species tree represents the best-supported topology based on species tree inference from multi-copy gene trees, determined using STAG, and rooted using STRIDE (Emms & Kelly, 2017). The genomes used for analysis included *Bombyx mori* (GCA_014905235.2), *Agriphila tristella* (GCA_928269145.1), *Micropterix aruncella* (GCA_944548615.1), *Papilio xuthus* (GCA_000836235.2), *Plutella xylostella* (GCA_932276165.1), *Achroia grisella* (GCA_030625045.1), *Acrobasis consociella* (GCA_963555685.1), *Acrobasis repandana* (GCA_963576875.1), *Acrobasis suavella* (GCA_943193695.1), *Amyelois transitella* (GCA_032362555.1), *Anerastia lotella* (GCA_964291865.1), *Apomyelois bistriatella* (GCA_947044815.1), *Cactoblastis cactorum* (GCA_020352625.1), *Corcyra cephalonica* (GCA_040436485.1), *Dioryctria mendacella* (GCA_964374295.1), *Elegia similella* (GCA_947532085.1), *Endotricha flammealis* (GCA_905163395.2), *Ephestia elutella* (GCA_018467065.1), *Ephestia kuehniella* (GCA_921024065.1), *Euzophera pinguis* (GCA_947363495.1), *Galleria mellonella* (GCA_026898425.1), *Homoeosoma sinuella* (GCA_964340625.1), *Hypsopygia costalis* (GCA_937001555.2), *Hypsopygia rubidalis* (GCA_965279855.1), *Pempelia palumbella* (GCA_964656235.1), *Phycita roborella* (GCA_964059365.1), *Plodia interpunctella* (GCA_027563975.2), *Pyralis farinalis* (GCA_947507595.1), *Pyralis regalis* (GCA_965194485.1), *Rhodophaea formosa* (GCA_963082605.1), *Synaphe punctalis* (GCA_965276045.1) and *Zophodia grossulariella* (GCA_965234215.1). The phylogenomic tree was generated using the packages ggtree v3.10.0 (Xu et al., 2022), aplot v0.2.2 (Yu, 2023), ape v5.7.1 (Paradis & Schliep, 2019) and tidyverse v2.0.0 (Wickham et al., 2019) in R v4.3.3 (R Core Team, 2024).

Synteny regions were determined using minimap2 v2.26 (Li, 2018), by aligning the assembled *A. sociella* genome to the publicly available genomes of *G. mellonella, A. grisella* and *C. cephalonica* using the parameter `-x asm10`. To generate synteny plots, the packages tidyverse v2.0.0, ggplot2 v3.5.2 (Wickham, 2016) and cowplot v1.1.3 (Wilke, 2024) were used in R v4.3.3.

## Results and discussion

### Genome assembly and annotation

Prior to this study, no publicly available genome assembly existed for *A. sociella*. We performed whole-genome sequencing, assembly, and annotation by isolating genomic DNA from a single *A. sociella* larvae. The genome sequence was assembled from 85.07 Gb data and 12.86 M ONT reads Four assemblies were generated using Canu with default settings and an adjusted error rate from 0.01 to 0.03. Assembly quality and completeness were quantitatively assessed based on evolutionarily informed expectations of gene content (**Fig. 1**). The Canu assembly generated with an adjusted error rate of 2% exhibited the most contiguous genome and was therefore selected for further refinement (**Table S1**).

**Fig. 1.**
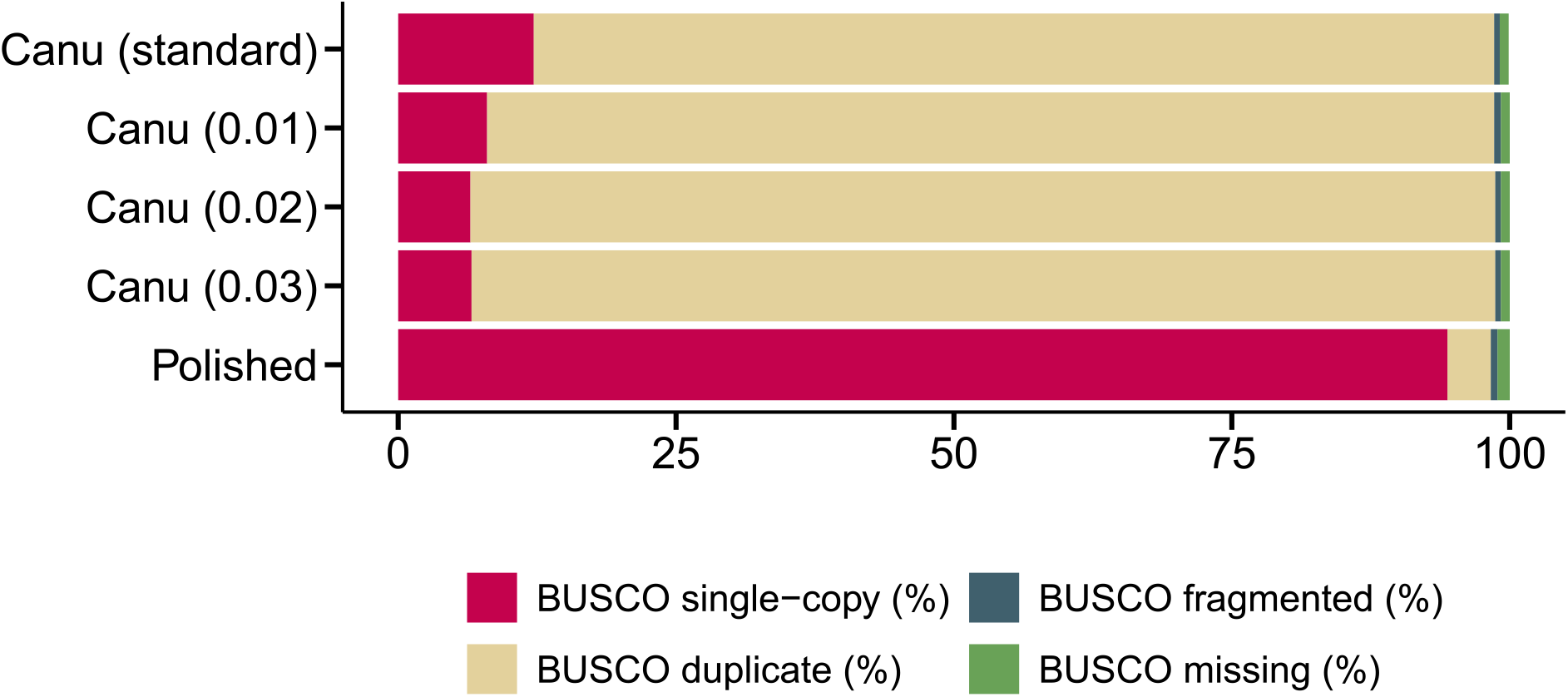
BUSCO scores of the Canu assemblies using standard parameters, and adjusted error rates of 1, 2 and 3%, as well as the polished assembly with reassigned haplotigs.

The selected assembly consisted of 1 294 Mb, 1 451 contigs, and N50 of 2.58 Mb while displaying a 92.2% duplication rate as determined by BUSCO (**Table S1**). Allelic contigs were reconciled and 47.6% (616 Mb) of the *A. sociella* genome was reassigned as haplotigs, and 0.22% (2.9 Mb) reassigned as artifacts. After polishing, the assembled genome had a total length of 675 Mb distributed across 347 contigs (N50 = 4.96 Mb). Based on the genome mode for BUSCO analysis, the assembly was 98.3% complete, 0.6% fragmented and had a duplication rate of 3.9% (**Fig. 1**). These metrics indicate that the genome assembly of *A. sociella* is of high quality.

RepeatModeler and RepeatMasker analyses classified and then masked 55.96% of the *A. sociella* genome for repetitive elements. Retroelements represented 33.40% of the genome, including Short Interspersed Nuclear Elements (14.90%), Long Interspersed Nuclear Elements (15.34%) and Long Terminal Repeats (3.16%). No satellite DNA was detected, and only 1.32% of the genome was represented by simple repeats. Retroelements were the most common categories of classified repeats, and some unclassified repeats (13.53%) were also detected.

Next, functional gene annotation for the genome assembly was obtained using a total of 28.1 Gb of data (28.3 M reads) from long-read RNA sequencing and 16.8 Gb of data (112.3 M reads) from short-read RNA sequencing. Annotation using Funannotate predicted a total of 42 009 genes, including 22 122 mRNAs and 20 140 tRNAs. 84.6% completeness was achieved using BUSCO completeness evaluation of genes, with 77.8% single copy genes and a duplication rate of 6.8% (**Table 2**). The inclusion long- and short-read RNAseq data reduced the total number of annotations compared to a workflow based on Funanntate and only short-read RNAseq data (50 287 genes, 27 242 mRNAs and 23 045 tRNAs; results not shown). Functional annotations with Pfam, InterPro and eggNOG identified at least one hit in 72.8% (16 105) of the genes, while 69.6% (15 394) of the genes were annotated as hypothetical proteins.

**Table 1.** Number of coding sequences, mRNAs, tRNAs and genes, and BUSCO completeness, duplications, and fragments for the annotated genome.

| Feature | Annotation |
| --- | --- |
| Coding Sequences (CDS) | 119 198 |
| Number of genes | 42 009 |
| Number of mRNAs | 22 122 |
| Number of tRNAs | 20 140 |
| **BUSCO on proteins** |  |
| BUSCO completeness (%) <sup>a</sup> | 84.6 |
| BUSCO single-copy (%) <sup>a</sup> | 77.8 |
| BUSCO duplicate (%) <sup>a</sup> | 6.8 |
| BUSCO fragmented (%) <sup>a</sup> | 8.6 |
| BUSCO missing (%) <sup>a</sup> | 6.8 |

<sup>a</sup> Determined using the BUSCO lepidoptera\_odb12 dataset.

### Phylogeny of *A. sociella*

A set of orthologs (BUSCO: lepidoptera_odb12 dataset) was chosen to perform phylogenomic analysis of the assembled genome of *A. sociella* and 31 species of the order Lepidoptera (**Table S1**). We recovered a robust phylogeny of the order Lepidoptera, with 28 genomes (including *A. sociella*) belonging to the family *Pyralidae*, with *B. mori* (family *Bombycidae*), *A. tristella* (family *Crambidae*), *M. aruncella* (family *Micropterigidae*), *P. xuthus* (family *Papilionidae*) and *P. xylostella* (family *Plutellidae*) included as outgroups. A*. sociella* is part of the subfamily *Galleriinae*, that also includes *G. mellonella, A. grisella* and *C. cephalonica* (**Fig. 1**). All nodes in the tree except one were supported with 100% bootstrap. *A. sociella* was placed in the same clade as *C. cephalonica* (rice moth), whose genus has been synonymized with *Aphomia* (Leraut, 2014). The phylogenomic relationship between these species reinforces the taxonomic classification. As expected, the closest relatives of *A. sociella*, correspond to species taxonomically classified as members of the *Galleriinae* subfamily (Scholtens & Solis, 2015).

Minimap2 was used to map the whole genome of *A. sociella* to its closest relatives in the *Galleriinae* subfamily. The syntenic blocks are generally less conserved between *A. sociella* and *G. mellonella* compared to *C. cephalonica* (**Fig. 2**). Nonetheless, multiple colinear blocks were identified across the genomes. Notably, the alignment between the genomes of *A. sociella* and *C. cephalonica* revealed longer and more continuous syntenic blocks between the genomes, indicating a higher degree of genome structure conservation.

**Fig. 2.**
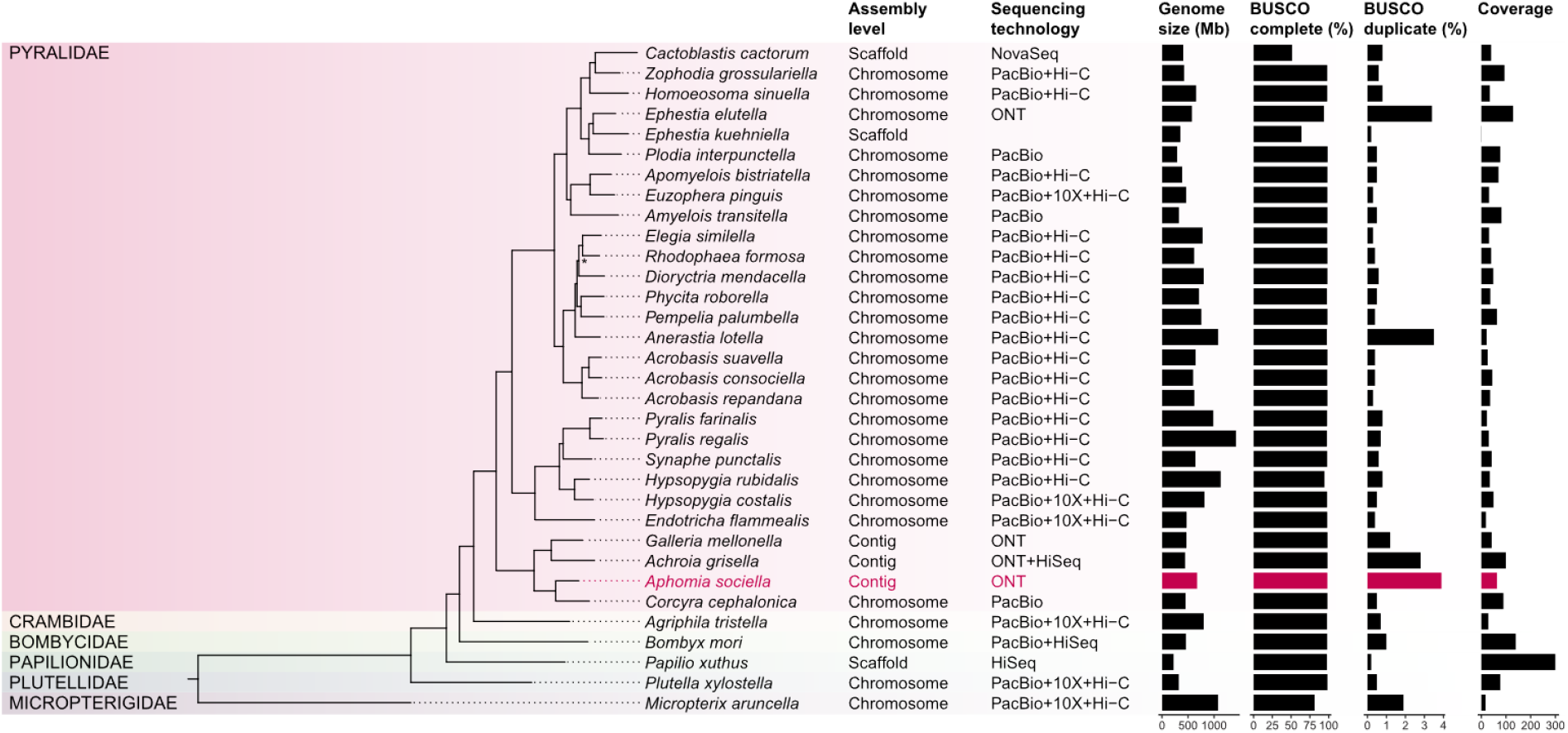
Phylogenetic placement of *A. sociella*. The tree was constructed using BUSCO single-copy orthologs (lepidoptera_odb12) from the assembled genome of this study along with publicly available genomes from other species of the Lepidoptera order. All nodes are supported by 100% bootstrap values, except one (indicated with asterisk, 61.1% bootstrap). Assembly level denotes the resolution of the genome, whether it is chromosome, scaffold of contig level. The sequencing platforms used for the assemblies are Illumina NovaSeq (NovaSeq), Illumina HiSeq (HiSeq), Pacific Biosciences (PacBio), Arima2 Hi-C (Hi-C), 10X Genomics Chromium (10X) and Oxford Nanopore Technologies (ONT), or a combination. The BUSCO complete (%) and BUSCO duplicate (%) are based on the lepidoptera_odb12 dataset.

**Fig. 3.**
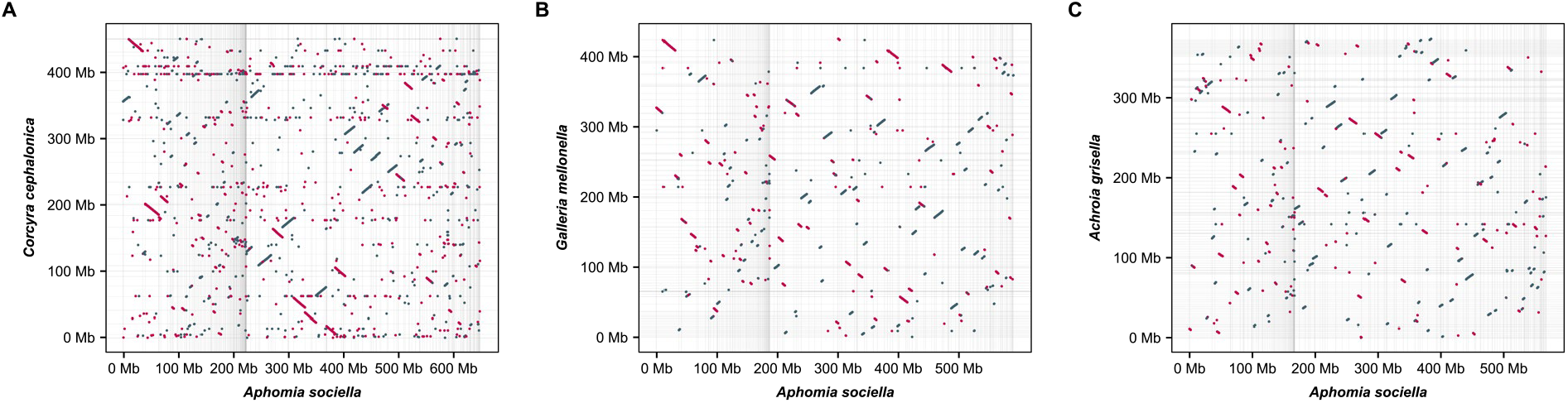
Synteny between the *A. sociella* genome and those of its closest relatives, **A)** *C. cephalonica* (GCA_040436485.1) **B)** *G. mellonella* (GCA_027563975.1) and **C)** *A. grisella* (GCA_030625045.1). Gridlines represent the boundary between contigs in the genome assemblies.

### Genes and pathways

The larvae of various species within the Lepidoptera order have frequently been observed consuming plastics, such as polyethylene (PE) and polystyrene. Indeed, larvae of *A. sociella* are also capable of chewing and ingesting PE films (**Fig.4**). Such behavior amongst larvae has attracted considerable scientific interest due to the potential use of these insects in the biodegradation and biotransformation of these recalcitrant synthetic polymers. Although *A. sociella* can penetrate through and consume plastic films, this behavior does not indicate an ability to biodegrade the plastic. Indeed, insects and their larvae eating through plastic film and packaging has long been a problem and has historically been the focus of prevention and control measures (Cline, 1978; Graham Bowditch, 1997; Highland, 1984; Sreenathan et al., 1960).

**Fig. 4.**
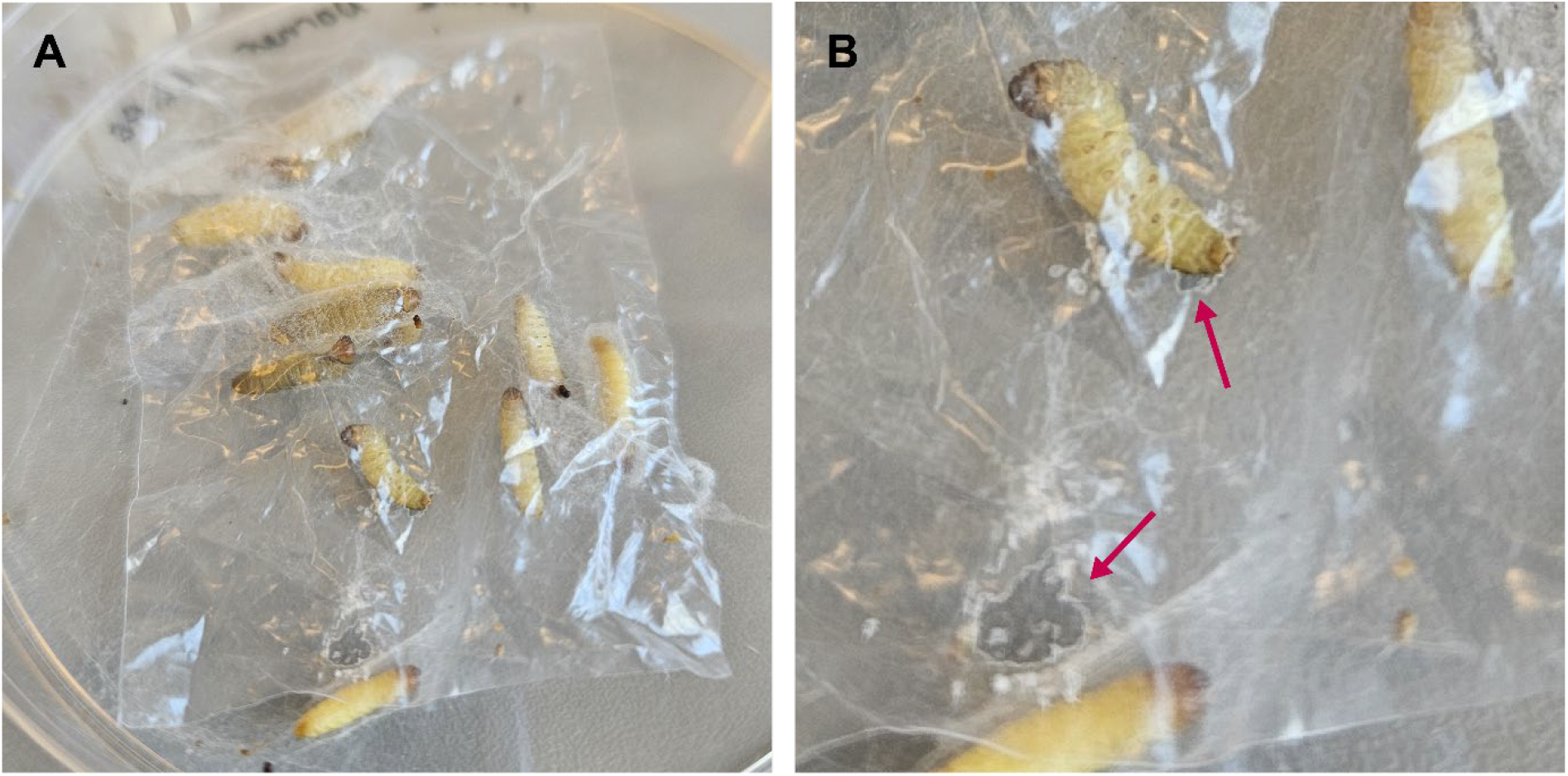
**A)** *A. sociella* larvae chewing on low-density polyethylene films. **B)** Holes in the polyethylene film made by the larvae.

Notably, in recent years, the larvae of *Galleria mellonella* have attracted considerable attention due to their ability to ingest PE (Peydaei et al., 2020; Sanluis-Verdes et al., 2022). Two proteins, an insect hexamerin and acidic juvenile-hormone suppressible protein, were proposed to confer *G. mellonella* the ability to oxidize this substrate (Sanluis-Verdes et al., 2022). However, their activity on PE could not be repeated in an independent study (Stepnov et al., 2024), which was in line with the findings of Réjasse et al (2022) who detected no bioassimilation of PE into the tissue of *G. mellonella* larvae when fed^2^H-labelled PE. (Réjasse et al., 2022). A more feasible approach to larval biotransformation of PE could entail a chemical pretreatment step to convert the relatively inert polymer into an oxy-functionalized, ow-molecular-weight waxy material that better resembles the natural wax metabolized by these insects. It has indeed been hypothesized that small oxidized aliphatic molecules derived from pretreated plastic can be further metabolized by the insect via the β-oxidation pathway (LeMoine et al., 2020). The genome of *A. sociella* contains genes encoding enzymes involved in the pathway for β-oxidation, including components responsible for the transport of long-chain fatty acids into the cytoplasm and mitochondria (**Table S2**). Multiple copies of cytochrome P450 (CYP) were also identified in the *A. sociella* genome. CYP is an oxidative enzyme suggested to be implicated in plastic degradation (Son et al., 2024), as it has been linked to the hydroxylation of hydrocarbon components of beeswax (Kong et al., 2019). It is important to note that the β-oxidation pathway and CYP have native functions beyond their potential involvement in plastic degradation, such as oxidizing lipids stored in the fat body of insect larvae (Gao et al., 2023), and oxidation/reduction of organic chemicals (Nauen et al., 2022). It is noteworthy that, for many lepidopteran species including pollinators, CYP have been shown to play a role in detoxification of secondary metabolites produced by plants and play a role in host adaptation (Heidel-Fischer & Vogel, 2015).

Pheromone signaling is an important aspect of the mating behavior of *A. sociella*, and the configuration of many sex pheromones has been characterized for this species (Wallin et al., 2020). The *A. sociella* genome contains genes encoding odorant-binding proteins (OBPs), chemosensory proteins (CSPs), odorant receptors (ORs) and sensory neuron membrane proteins (SNMPs) (**Table S4**), which are most likely involved in the pheromone signaling between the moths (Jiang et al., 2021; Pregitzer et al., 2014). Understanding the underlying mechanisms involved in pheromone synthesis and olfaction can serve as a way to disrupt mating, thus acting as a pest control measure (Kong et al., 2014).

Toll-like receptors (TLRs) are essential components of signaling pathways that regulate both developmental processes and innate immune responses in insects. The *A. sociella* genome encodes multiple TLRs, including proteins predicted to contain both toll/interleukin-1 receptor homology (TIR) domains and leucine-rich repeat (LRR) domains. Insect TLRs are typically characterized by the presence of TIRs and LRRs (Imler & Zheng, 2003), and all the 14 putative TLR proteins contain his conserved domain architecture (**Table S3**). Strategies to control moth population, for instance the gypsy moth (*Lymantria dispar*), include the use of “bioinsecticides” (Zhang et al., 2019), which are designed to disrupt the immune systems of the insect. Therefore, characterizing the presence and structure of TLRs in the bumblebee wax moth is essential for informing the development of effective bioinsecticide-based strategies.

## Conclusion

Here, we present a high-quality contig-level genome assembly of *A. sociella*. A combination of short- and long-read transcriptomic data was utilized to achieve comprehensive and accurate functional gene annotation. This enabled the identification of genes involved in fundamental biological processes as well as those associated with species-specific behaviors, providing valuable insights into the molecular basis of *A. sociella* biology. As expected, phylogenomic analysis placed *A. sociella* in the same clade as *C. cephalonica* and indicated close evolutionary relationships with the other two species in the subfamily *Galleriinae*, namely *A. grisella* and *G. mellonella*. We expect that the availability of an annotated genome for *A. sociella* will facilitate further research into the life cycle and behavior of the moth and can be used in future research on pest control measures. In addition, the genome sequence will facilitate microbiome studies by enabling the effective removal of contaminating host DNA in metagenomic analyses.

## Data availability

The Nanopore whole genome sequencing raw data are available through the Sequence Read Archive (SRA) with the accession number PRJNA1276471. The genome assembly is available through the NCBI GenBank database with the accession number SAMN49073243. The RNA sequencing reads are available through the SRA with the following accession numbers: SAMN49327582 (Illumina short-read RNAseq) and SAMN49327583 (Nanopore long-read RNAseq), both used to annotate the genome assembly. Gene annotations have been deposited in Figshare (DOI: 10.6084/m9.figshare.29279816).

## Acknowledgments

This work was supported by the Norwegian University of Life Sciences through the Sustainability Arena program SmartPlast, and the Research Council of Norway under Grant no. 326975 (Enzyclic). We thank Atle Mjelde for his contributions in supplying us with *A. sociella* larvae, as well as providing us with his knowledge of this species. We acknowledge the Orion High Performance Computing Centre and the Threadripper station at the Norwegian University of Life Sciences, for computational resources that contributed to producing the research results reported within this paper. The authors thank Elixir-Norway for bioinformatics and data management related services.

## Supplementary data

**Table S1:** Statistics for Canu assemblies and the polished genome.

**Table S2:** Available reference genomes for species in the order Lepidoptera used for phylogenomic analysis. The species marked in red are from this study (excel file).

**Table S3:** Genes hypothesized to be involved in plastic biodegradation.

**Table S4:** Immune genes.

**Table S5:** Pheromone genes.

## References

Andrews, S. (2010). FastQC: A Quality Control Tool for High Throughput Sequence Data. In http://www.bioinformatics.babraham.ac.uk/projects/fastqc/

Aramaki, T., Blanc-Mathieu, R., Endo, H., Ohkubo, K., Kanehisa, M., Goto, S., & Ogata, H. (2020). KofamKOALA: KEGG Ortholog assignment based on profile HMM and adaptive score threshold. Bioinformatics, 36(7), 2251–2252. 10.1093/bioinformatics/btz859

Ashburner, M., Ball, C. A., Blake, J. A., Botstein, D., Butler, H., Cherry, J. M., Davis, A. P., Dolinski, K., Dwight, S. S., Eppig, J. T., Harris, M. A., Hill, D. P., Issel-Tarver, L., Kasarskis, A., Lewis, S., Matese, J. C., Richardson, J. E., Ringwald, M., Rubin, G. M., & Sherlock, G. (2000). Gene Ontology: tool for the unification of biology. Nature Genetics, 25(1), 25–29. 10.1038/75556

Astashyn, A., Tvedte, E. S., Sweeney, D., Sapojnikov, V., Bouk, N., Joukov, V., Mozes, E., Strope, P. K., Sylla, P. M., Wagner, L., Bidwell, S. L., Clark, K., Davis, E. W., Smith-White, B., Hlavina, W., Pruitt, K. D., Schneider, V. A., & Murphy, T. D. (2023). Rapid and sensitive detection of genome contamination at scale with FCS-GX. Cold Spring Harbor Laboratory. 10.1101/2023.06.02.543519

Bushmanova, E., Antipov, D., Lapidus, A., & Prjibelski, A. D. (2019). rnaSPAdes: a de novo transcriptome assembler and its application to RNA-Seq data. GigaScience, 8(9). 10.1093/gigascience/giz100

Cantalapiedra, C. P., Hernández-Plaza, A., Letunic, I., Bork, P., & Huerta-Cepas, J. (2021). eggNOG-mapper v2: Functional Annotation, Orthology Assignments, and Domain Prediction at the Metagenomic Scale. Molecular Biology and Evolution, 38(12), 5825–5829. 10.1093/molbev/msab293

Chen, S., Zhou, Y., Chen, Y., & Gu, J. (2018). fastp: an ultra-fast all-in-one FASTQ preprocessor. Bioinformatics, 34(17), i884–i890. 10.1093/bioinformatics/bty560

Chen, Y., Zhang, Y., Wang, A. Y., Gao, M., & Chong, Z. (2021). Accurate long-read de novo assembly evaluation with Inspector. Genome Biology, 22(1). 10.1186/s13059-021-02527-4

Cline, D. L. (1978). Penetration of Seven Common Flexible Packaging Materials by Larvae and Adults of Eleven Species of Stored-Product Insects. Journal of Economic Entomology, 71(5), 726–729. 10.1093/jee/71.5.726

Consortium, T. G. O., Aleksander, S. A., Balhoff, J., Carbon, S., Cherry, J. M., Drabkin, H. J., Ebert, D., Feuermann, M., Gaudet, P., Harris, N. L., Hill, D. P., Lee, R., Mi, H., Moxon, S., Mungall, C. J., Muruganugan, A., Mushayahama, T., Sternberg, P. W., Thomas, P. D., … Westerfield, M. (2023). The Gene Ontology knowledgebase in 2023. Genetics, 224(1). 10.1093/genetics/iyad031

Consortium, T. U. (2022). UniProt: the Universal Protein Knowledgebase in 2023. Nucleic Acids Research, 51(D1), D523–D531. 10.1093/nar/gkac1052

Drula, E., Garron, M.-L., Dogan, S., Lombard, V., Henrissat, B., & Terrapon, N. (2021). The carbohydrate-active enzyme database: functions and literature. Nucleic Acids Research, 50(D1), D571–D577. 10.1093/nar/gkab1045

Emms, D. M., & Kelly, S. (2017). STRIDE: Species Tree Root Inference from Gene Duplication Events. Molecular Biology and Evolution, 34(12), 3267–3278. 10.1093/molbev/msx259

Emms, D. M., & Kelly, S. (2019). OrthoFinder: phylogenetic orthology inference for comparative genomics. Genome Biology, 20(1). 10.1186/s13059-019-1832-y

Gao, X., Zhang, J., Qin, Q., Wu, P., Zhang, H., & Meng, Q. (2023). Metabolic changes during larval– pupal metamorphosis of Helicoverpa armigera. Insect Science, 30(6), 1663–1676. 10.1111/1744-7917.13201

Graham Bowditch, T. (1997). Penetration of Polyvinyl Chloride and Polypropylene Packaging Films by Ephestia cautella (Lepidoptera: Pyralidae) and Plodia interpunctella (Lepidoptera: Pyralidae) Larvae, and Tribolium confusum (Coleoptera: Tenebrionidae) Adults. Journal of Economic Entomology, 90(4), 1028–1031. 10.1093/jee/90.4.1028

Heidel-Fischer, H. M., & Vogel, H. (2015). Molecular mechanisms of insect adaptation to plant secondary compounds. Current Opinion in Insect Science, 8, 8–14. 10.1016/j.cois.2015.02.004

Highland, H. A. (1984). Insect infestation of packages. Insect management for food storage and processing, 309–320.

Imler, J.-L., & Zheng, L. (2003). Biology of Toll receptors: lessons from insects and mammals. Journal of Leukocyte Biology, 75(1), 18–26. 10.1189/jlb.0403160

Jiang, X.-C., Liu, S., Jiang, X.-Y., Wang, Z.-W., Xiao, J.-J., Gao, Q., Sheng, C.-W., Shi, T.-F., Zeng, H.-R., Yu, L.-S., & Cao, H.-Q. (2021). Identification of Olfactory Genes From the Greater Wax Moth by Antennal Transcriptome Analysis. Frontiers in Physiology, 12. 10.3389/fphys.2021.663040

Jones, P., Binns, D., Chang, H.-Y., Fraser, M., Li, W., McAnulla, C., McWilliam, H., Maslen, J., Mitchell, A., Nuka, G., Pesseat, S., Quinn, A. F., Sangrador-Vegas, A., Scheremetjew, M., Yong, S.-Y., Lopez, R., & Hunter, S. (2014). InterProScan 5: genome-scale protein function classification. Bioinformatics, 30(9), 1236–1240. 10.1093/bioinformatics/btu031

Kanehisa, M., & Sato, Y. (2020). KEGG Mapper for inferring cellular functions from protein sequences. Protein Science, 29(1), 28–35. 10.1002/pro.3711

Khramov, P. (2025). Illustrated Insecta catalogue. https://insecta.pro/

Kindl, J., Jiroš, P., Kalinová, B., Žáček, P., & Valterová, I. (2012). Females of the Bumblebee Parasite, Aphomia sociella, Excite Males Using a Courtship Pheromone. Journal of Chemical Ecology, 38(4), 400–407. 10.1007/s10886-012-0100-3

Kindl, J., Kalinová, B., Červenka, M., Jílek, M., & Valterová, I. (2011). Male Moth Songs Tempt Females to Accept Mating: The Role of Acoustic and Pheromonal Communication in the Reproductive Behaviour of Aphomia sociella. PLoS ONE, 6(10), e26476. 10.1371/journal.pone.0026476

Kong, H. G., Kim, H. H., Chung, J.-H., Jun, J., Lee, S., Kim, H.-M., Jeon, S., Park, S. G., Bhak, J., & Ryu, C.-M. (2019). The Galleria mellonella Hologenome Supports Microbiota-Independent Metabolism of Long-Chain Hydrocarbon Beeswax. Cell Reports, 26(9), 2451–2464.e2455. 10.1016/j.celrep.2019.02.018

Kong, W. N., Li, J., Fan, R. J., Li, S. C., & Ma, R. Y. (2014). Sex-Pheromone-Mediated Mating Disruption Technology for the Oriental Fruit Moth, Grapholita molesta (Busck) (Lepidoptera: Tortricidae): Overview and Prospects. Psyche: A Journal of Entomology, 2014, 1–8. 10.1155/2014/253924

Koren, S., Walenz, B. P., Berlin, K., Miller, J. R., Bergman, N. H., & Phillippy, A. M. (2017). Canu: scalable and accurate long-read assembly via adaptive k-mer weighting and repeat separation. Genome Research, 27(5), 722–736. 10.1101/gr.215087.116

Krams, R., Grigorjeva, T., Willow, J., Popovs, S., Munkevics, M., Trakimas, G., Contreras-Garduño, J., De Souza, A. R., Adams, C. B., Rantala, M. J., Garajeva, S. J., Sledevskis, E., Krama, T., & Krams, I. A. (2025). Infestation levels of Aphomia sociella in bumblebees increase with proximity to apiaries and result in lower reproductive output and weaker immune response. Frontiers in Bee Science, 3. 10.3389/frbee.2025.1550560

LeMoine, C. M. R., Grove, H. C., Smith, C. M., & Cassone, B. J. (2020). A Very Hungry Caterpillar: Polyethylene Metabolism and Lipid Homeostasis in Larvae of the Greater Wax Moth (Galleria mellonella). Environmental Science & Technology, 54(22), 14706–14715. 10.1021/acs.est.0c04386

Leraut, P. (2014). Moths of Europe, Volume 4, Pyralids 2. NAP Editions.

Lewin, H. A., Richards, S., Lieberman Aiden, E., Allende, M. L., Archibald, J. M., Bálint, M., Barker, K. B., Baumgartner, B., Belov, K., Bertorelle, G., Blaxter, M. L., Cai, J., Caperello, N. D., Carlson, K., Castilla-Rubio, J. C., Chaw, S.-M., Chen, L., Childers, A. K., Coddington, J. A., … Zhang, G. (2022). The Earth BioGenome Project 2020: Starting the clock. Proceedings of the National Academy of Sciences, 119(4), e2115635118. 10.1073/pnas.2115635118

Li, H. (2018). Minimap2: pairwise alignment for nucleotide sequences. Bioinformatics, 34(18), 3094–3100. 10.1093/bioinformatics/bty191

Manni, M., Berkeley, M. R., Seppey, M., & Zdobnov, E. M. (2021). BUSCO: Assessing Genomic Data Quality and Beyond. Current Protocols, 1(12). 10.1002/cpz1.323

Mistry, J., Chuguransky, S., Williams, L., Qureshi, M., Salazar Gustavo A., Sonnhammer, E. L. L., Tosatto, S. C. E., Paladin, L., Raj, S., Richardson, L. J., Finn, R. D., & Bateman, A. (2020). Pfam: The protein families database in 2021. Nucleic Acids Research, 49(D1), D412–D419. 10.1093/nar/gkaa913

Mugford, S., Wouters, R., Mathers, T. C., & Hogenhout, S. (2020). High quality DNA extraction from very small individual insects. protocols.io

Nauen, R., Bass, C., Feyereisen, R., & Vontas, J. (2022). The Role of Cytochrome P450s in Insect Toxicology and Resistance. Annual Review of Entomology, 67(1), 105–124. 10.1146/annurev-ento-070621-061328

Nielsen, H., Tsirigos, K. D., Brunak, S., & Von Heijne, G. (2019). A Brief History of Protein Sorting Prediction. The Protein Journal, 38(3), 200–216. 10.1007/s10930-019-09838-3

Niu, J., Chen, R., & Wang, J. J. (2024). RNA interference in insects: the link between antiviral defense and pest control. Insect Science, 31(1), 2–12. 10.1111/1744-7917.13208

Oxford Nanopore Technologies. (2023). Guppy protocol. Retrieved July 11th from https://nanoporetech.com/document/Guppy-protocol

Palmer, J. (2016). Funannotate: pipeline for genome annotation. In https://zenodo.org/records/2604804

Paradis, E., & Schliep, K. (2019). ape 5.0: an environment for modern phylogenetics and evolutionary analyses in R. Bioinformatics, 35(3), 526–528. 10.1093/bioinformatics/bty633

Peydaei, A., Bagheri, H., Gurevich, L., de Jonge, N., & Nielsen, J. L. (2020). Impact of polyethylene on salivary glands proteome in Galleria melonella. Comparative Biochemistry and Physiology Part D: Genomics and Proteomics, 34, 100678. 10.1016/j.cbd.2020.100678

Pregitzer, P., Greschista, M., Breer, H., & Krieger, J. (2014). The sensory neurone membrane protein SNMP1 contributes to the sensitivity of a pheromone detection system. Insect Molecular Biology, 23(6), 733–742. 10.1111/imb.12119

R Core Team. (2024). R: A Language and Environment for Statistical Computing. In https://www.R-project.org/

Rawlings, N. D., Barrett, A. J., Thomas, P. D., Huang, X., Bateman, A., & Finn, R. D. (2017). The MEROPS database of proteolytic enzymes, their substrates and inhibitors in 2017 and a comparison with peptidases in the PANTHER database. Nucleic Acids Research, 46(D1), D624–D632. 10.1093/nar/gkx1134

Réjasse, A., Waeytens, J., Deniset-Besseau, A., Crapart, N., Nielsen-Leroux, C., & Sandt, C. (2022). Plastic biodegradation: Do Galleria mellonella Larvae Bioassimilate Polyethylene? A Spectral Histology Approach Using Isotopic Labeling and Infrared Microspectroscopy. Environmental Science &amp; Technology, 56(1), 525–534. 10.1021/acs.est.1c03417

Ren, L., Men, L., Zhang, Z., Guan, F., Tian, J., Wang, B., Wang, J., Zhang, Y., & Zhang, W. (2019). Biodegradation of Polyethylene by Enterobacter sp. D1 from the Guts of Wax Moth Galleria mellonella. International Journal of Environmental Research and Public Health, 16(11), 1941. 10.3390/ijerph16111941

Rhie, A., Walenz, B. P., Koren, S., & Phillippy, A. M. (2020). Merqury: reference-free quality, completeness, and phasing assessment for genome assemblies. Genome Biology, 21(1). 10.1186/s13059-020-02134-9

Roach, M. J., Schmidt, S. A., & Borneman, A. R. (2018). Purge Haplotigs: allelic contig reassignment for third-gen diploid genome assemblies. BMC Bioinformatics, 19(1). 10.1186/s12859-018-2485-7

Sanluis-Verdes, A., Colomer-Vidal, P., Rodriguez-Ventura, F., Bello-Villarino, M., Spinola-Amilibia, M., Ruiz-Lopez, E., Illanes-Vicioso, R., Castroviejo, P., Aiese Cigliano, R., Montoya, M., Falabella, P., Pesquera, C., Gonzalez-Legarreta, L., Arias-Palomo, E., Solà, M., Torroba, T., Arias, C. F., & Bertocchini, F. (2022). Wax worm saliva and the enzymes therein are the key to polyethylene degradation by Galleria mellonella. Nature Communications, 13(1). 10.1038/s41467-022-33127-w

Schoch, C. L., Ciufo, S., Domrachev, M., Hotton, C. L., Kannan, S., Khovanskaya, R., Leipe, D., McVeigh, R., O’Neill, K., Robbertse, B., Sharma, S., Soussov, V., Sullivan, J. P., Sun, L., Turner, S., & Karsch-Mizrachi, I. (2020). NCBI Taxonomy: a comprehensive update on curation, resources and tools. Database (Oxford), 2020. 10.1093/database/baaa062

Scholtens, B., & Solis, M. A. (2015). Annotated check list of the Pyraloidea (Lepidoptera) of America North of Mexico. ZooKeys, 535, 1–136. 10.3897/zookeys.535.6086

Smit, A., & Hubley, R. (2008-2015). RepeatModeler Open-1.0. In http://www.repeatmasker.org

Smit, A., Hubley, R., & Green, P. (2013-2015). RepeatMasker Open-4.0. In http://www.repeatmasker.org

Son, J.-S., Lee, S., Hwang, S., Jeong, J., Jang, S., Gong, J., Choi, J. Y., Je, Y. H., & Ryu, C.-M. (2024). Enzymatic oxidation of polyethylene by Galleria mellonella intestinal cytochrome P450s. Journal of Hazardous Materials, 480, 136264. 10.1016/j.jhazmat.2024.136264

Sreenathan, V. R., Iyengar, N. V. R., Narasimhan, K. S., & Majumder, S. K. (1960). Studies on insect resistance of packaging materials-cellulose and polyethylene films. Food Science, 9(6), 3.

Stepnov, A. A., Lopez-Tavera, E., Klauer, R., Lincoln, C. L., Chowreddy, R. R., Beckham, G. T., Eijsink, V. G. H., Solomon, K., Blenner, M., & Vaaje-Kolstad, G. (2024). Revisiting the activity of two poly(vinyl chloride)- and polyethylene-degrading enzymes. Nature Communications, 15(1). 10.1038/s41467-024-52665-z

Trizna, M. (2020). assembly_stats. Zenodo. 10.5281/zenodo.3968775

van Nieukerken, E. J., Kaila, L., Kitching, I., Kristensen, N. P., Lees, D., Minet, J., Mitter, C., Mutanen, M., Regier, J. C., Simonsen, T., Wahlberg, N., Yen, S.-H., Zahiri, R., Adamski, D., Baixeras, J., Bartsch, D., Bengtsson, B. Å., Brown, J. W., Bucheli, S., & Zwick, A. (2011). Order Lepidoptera Linnaeus, 1758. In Z. Z-Q (Ed.), Animal Biodiversity: An outline of higher classification and survey of taxonomic richness (Vol. 3148, pp. 212–221). Zootaxa.

Wallin, E. A., Kalinová, B., Kindl, J., Hedenström, E., & Valterová, I. (2020). Stereochemistry of two pheromonal components of the bumblebee wax moth, Aphomia sociella. Scientific Reports, 10(1). 10.1038/s41598-020-59069-1

Wickham, H. (2016). ggplot2: Elegant Graphics for Data Analysis. Springer-Verlag. https://ggplot2.tidyverse.org

Wickham, H., Averick, M., Bryan, J., Chang, W., McGowan, L. D. A., François, R., Grolemund, G., Hayes, A., Henry, L., Hester, J., Kuhn, M., Pedersen, T. L., Miller, E., Bache, S. M., Müller, K., Ooms, J., Robinson, D., Seidel, D. P., Spinu, V., … Yutani, H. (2019). Welcome to the tidyverse. Journal of Open Source Software, 4(43). 10.21105/joss.01686

Wilke, C. O. (2024). cowplot: Streamlined Plot Theme and Plot Annotations for ‘ggplot2’. In https://CRAN.R-project.org/package=cowplot

Witzgall, P., Kirsch, P., & Cork, A. (2010). Sex Pheromones and Their Impact on Pest Management. Journal of Chemical Ecology, 36(1), 80–100. 10.1007/s10886-009-9737-y

Xu, S., Li, L., Luo, X., Chen, M., Tang, W., Zhan, L., Dai, Z., Lam, T. T., Guan, Y., & Yu, G. (2022). Ggtree: A serialized data object for visualization of a phylogenetic tree and annotation data. iMeta, 1(4). 10.1002/imt2.56

Yu, G. (2023). aplot: Decorate a ‘ggplot’ with Associated Information. In https://CRAN.R-project.org/package=aplot

Zhang, J., Cong, Q., Rex, E. A., Hallwachs, W., Janzen, D. H., Grishin, N. V., & Gammon, D. B. (2019). Gypsy moth genome provides insights into flight capability and virus–host interactions. Proceedings of the National Academy of Sciences, 116(5), 1669–1678. 10.1073/pnas.1818283116

Zhang, J., Gao, D., Li, Q., Zhao, Y., Li, L., Lin, H., Bi, Q., & Zhao, Y. (2020). Biodegradation of polyethylene microplastic particles by the fungus Aspergillus flavus from the guts of wax moth Galleria mellonella. Science of the Total Environment, 704, 135931. 10.1016/j.scitotenv.2019.135931

Zhang, Q., Dou, W., Taning, C. N. T., Smagghe, G., & Wang, J.-J. (2021). Regulatory roles of microRNAs in insect pests: prospective targets for insect pest control. Current Opinion in Biotechnology, 70, 158–166. 10.1016/j.copbio.2021.05.002

